# Crop loss identification at field parcel scale using satellite remote sensing and machine learning

**DOI:** 10.1101/2021.05.07.443072

**Authors:** Santosh Hiremath, Samantha Wittke, Taru Palosuo, Jere Kaivosoja, Fulu Tao, Maximilian Proll, Eetu Puttonen, Pirjo Peltonen-Sainio, Pekka Marttinen, Hiroshi Mamitsuka

**Author notes:** These two authors contributed equally to this work (Santosh Hiremath, Samantha Wittke). These three authors contributed equally to this work (Pirjo Peltonen-Sainio, Pekka Marttinen, Hiroshi Mamitsuka).

## Abstract

Identifying crop loss at field parcel scale using satellite images is challenging: first, crop loss is caused by many factors during the growing season; second, reliable reference data about crop loss are lacking; third, there are many ways to define crop loss. This study investigates the feasibility of using satellite images to train machine learning (ML) models to classify agricultural field parcels into those with and without crop loss. The reference data for this study was provided by Finnish Food Authority (FFA) containing crop loss information of approximately 1.4 million field parcels in Finland covering about 3.5 million ha from 2000 to 2015. This reference data was combined with Normalised Difference Vegetation Index (NDVI) derived from Landsat 7 images, in which more than 80% of the possible data are missing. Despite the hard problem with extremely noisy data, among the four ML models we tested, random forest (with mean imputation and missing value indicators) achieved the average AUC (area under the ROC curve) of 0.688 *±* 0.059 over all 16 years with the range [0.602, 0.795] in identifying new crop-loss fields based on reference fields of the same year. To our knowledge, this is one of the first large scale benchmark study of using machine learning for crop loss classification at field parcel scale. The classification setting and trained models have numerous potential applications, for example, allowing government agencies or insurance companies to verify crop-loss claims by farmers and realise efficient agricultural monitoring.

## 1 Introduction

Future food production is challenged by increasing demand for more sustainable agricultural systems that consider environmental, economic and social dimensions of sustainability. Remote sensing data from spaceborne platforms offer a possibility to address these challenges [1]. Multispectral satellite remote sensing applications in agriculture date back to the early 1970s with the launch of Landsat 1 by the National Aeronautics and Space Agency (NASA). Applications include agriculture land use mapping [2], agricultural monitoring [3], leaf area index (LAI) and biomass estimation [4, 5], precision agriculture [6], agricultural water management [7], estimation of crop yield [8–13], and crop damage assessment caused by floods [14, 15] and lodging [16]. These studies have focused mostly on the regional scale due to a spatially limited reference data at a field parcel scale.

Only a few studies have considered the use of remote sensing data from spaceborne platforms to assess crop loss at field parcel scale. For example, multispectral spaceborne remote sensing data from Landsat ETM+ was used to study the effect of sowing date and weed control during fallow period on spring wheat yield in Mexico by [17]. Based on experiments on 100 fields across three seasons they concluded that the effect of sowing data and weed control on yield can be estimated using multispectral spaceborne remote sensing data. Tapia-Silva et al. [18] studied crop losses due to flood using Landsat TM/ETM on 132 field parcels across 15 seasons and concluded that modelling crop loss was challenging. Data from Sentinel-1 and 2 satellites were used by [19] for cyclone damage assessment on 200 coconut and 200 rice fields in India with promising results. Crop damages on 600 wheat fields were also studied by [20] in Greece using spaceborne multispectral imagery and ancillary geospatial data, but they faced the challenge of defining field parcels based on low resolution imagery available from spaceborne platforms. Two recent studies [21, 22] used Synthetic Aperture Radar (SAR) images to assess crop damage due the 2020 wind storm Derecho. They apply their method across the state of Iowa, United States to estimate the area of crop damage and verify their methods with ground truth data of a maximum of 14 fields. A common theme across many of the above studies is the limited number of ground truth data from field parcels used to evaluate the proposed methods. This is understandable because collecting large amounts of ground truth data from many field parcels is challenging. In summary, to the best of our knowledge, remote sensing applications have been limited to specific crop-loss type based on its cause such as flooding or drought and are also limited to coarse spatial scale due to lack of ground truth data. A large study about the use of remote sensing data for monitoring general crop loss (not specific to any causal factor) at field parcel scale is desirable. Because it can aid both public and private organisations like insurance companies and governments, respectively, in agricultural monitoring [23, 24]. Such a study can also aid in effective implementation of Common Agricultural Policy (CAP) reforms enacted by the European Commission (EC) [25]. In fact EC has started projects such as Sentinels for CAP (Sen4Cap), aimed at developing new remote sensing methods for agricultural monitoring at scale [26].

There are many drivers for crop loss. Yield potential in a given field and region depends on the crop and cultivar. In addition to a soil type and weather conditions, farmer’s decisions have an impact on yield potential and the risk for crop loss. The risk of crop loss can be reduced by using quality seeds [27], planting well-adapted cultivars [28], matching crops to the most appropriate field parcels [29], as well as by adopting timely and accurate management practices such as sowing, crop protection and harvesting. For high-latitude agricultural systems risks caused by variable weather are substantial, and total large scale crop failures may occur once or twice a decade [30]. In this study, we use the definition of crop loss of Finnish crop damage compensation program. It is given by the percentage of area of field parcel. This definition decouples crop loss from its driver allowing the study of general crop loss irrespective of the driver.

This study aims to test the feasibility of combining machine learning (ML) models with optical satellite data to classify field parcels with and without crop loss. The reference data used consists of approximately 1.4 million barley (*Hordeum vulgare* L.) fields covering 3.5 million ha in Finland. The time period of the study is sixteen years from 2000 to 2015. Two settings are considered: 1) within-year classification, where training and test data are from the same year and 2) between-year classification, where training and test data are from different years. Both can be applied to the task of verifying a crop loss reported by a farmer, while 1) corresponds to the situation where data from other fields are available in the respective year, whereas in 2) no such data are available. The overview of the study is illustrated in the form of a flowchart in Fig 1. Performance results (AUC) obtained for within-year classification was approximately 0.7 on average over 16 years, while the regression line estimated by the projection of our results implied this performance can be improved if the missing data ratio was reduced. Analysis of the results revealed high amount of missing data in satellite image time series (more than 80% in our case) can have a significant impact on the classification performance. Thanks to the very comprehensive data set and wide spread of the area of investigation, we expect that our conclusions regarding the classification performance of the methods to be robust and generalisable for Barley in other countries.

**Fig 1.**
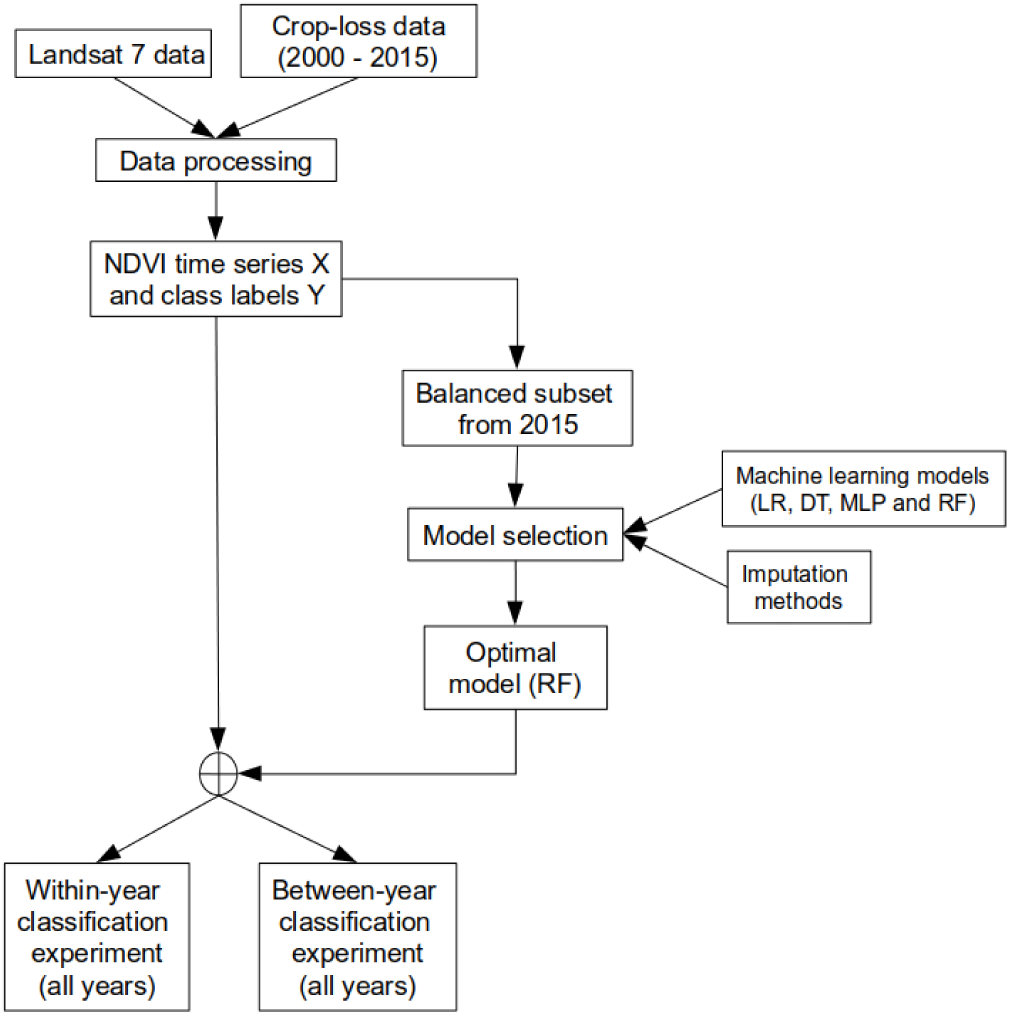
Overview of the study as a flowchart.

## 2 Materials and methods

### 2.1 Study area and crop loss data

The study area includes southern and western regions of Finland from 2000 to 2015. The area investigated covers the coastal agricultural land area in Finland comprising of 1.4 million field parcels growing barley and covering approximately 3.5 million ha. The size of the field parcels varied from 1 to 90 ha with an average of 2.4 ha. The study area and the distribution of the field parcels are shown in Fig 2.

**Fig 2.**
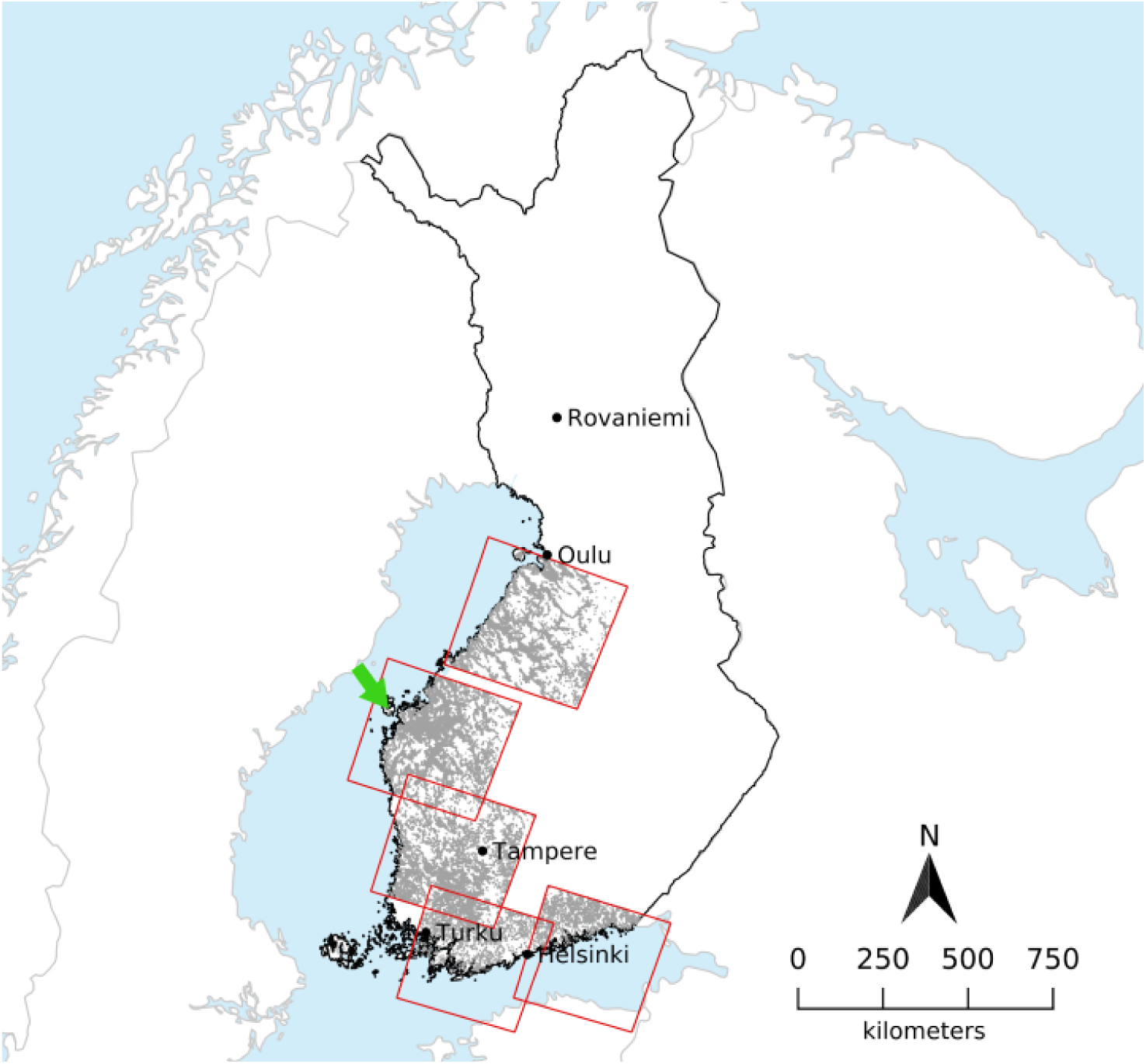
Study area. Landsat-7 tiles extent (red) over South-Western Finland utilised in this study with barley fields of the year 2000 added in grey. Centre coordinates (latitude, longitude) of the Landsat 7 tiles from North to South: 19015: 64.22478116, 25.13267424; 19116: 62.85736322, 22.45274870 ; 19017: 61.48156017, 22.95909640; 18918 (southwestern): 60.10043047, 23.54123385; 18718 (southeastern): 60.09748833, 26.63623269; background: Natural Earth, Finland: National Land Survey of Finland Topographic Database 05/2021. The green arrow denotes the location of the subset shown in Figure 3

The reference data on crop loss was provided by Finnish Food Authority (FFA). The data consisted of field parcel ID, field boundary (see example area in Figure 3), area, crop type and variety, crop loss (as area of the field parcel) and farm ID for the years from 2000 to 2015. The data originates from the crop damage compensation system in Finland that started in 1976 and lasted until 2015 [31]. The analysis was made with barley as it most amount of reference data among all the crops.

**Fig 3.**
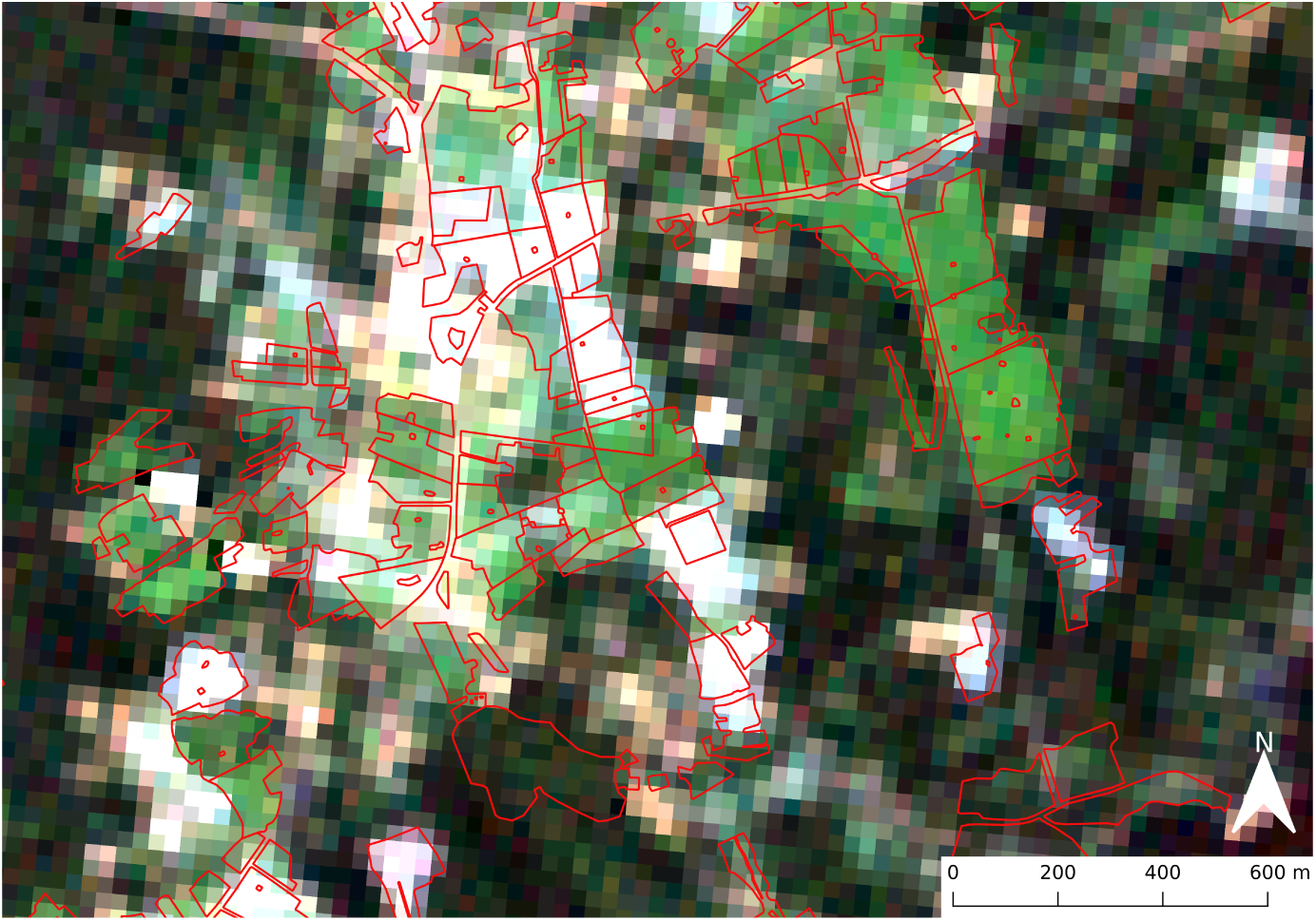
Field parcel example. Example area with field parcels boundaries (red) from the year 2012 (open data: http://www.nic.funet.fi/index/geodata/mavi/peltolohkot/2012/, accessed 28.09.21) and Landsat 7 true color image from 03.08.01 in the background (location: Western Finland, green arrow in Figure 2)

The crop loss data were collected through a self-reporting survey where the farmers reported crop loss as percentage of the area of the field. This was processed into a binary variable where anything greater than zero percent indicated crop loss (1) and everything else as indicated no loss (0). The number of field parcels with and without crop loss for each year from 2000 to 2015 is shown in Table 1. Over all years and the whole area of investigation, there were 33,840 field parcels (2.38%) with crop loss and 1,418,872 (97.62%) with no loss. We observed that larger fields had reported loss more often than smaller fields so the effect of field size on crop loss classification performance was examined (see the last part of Section 3.2).

**Table 1.** Percentage of field parcels for which crop loss was reported for each year.

| year | #parcels | #parcels<br>with loss | loss ratio (%) |
| --- | --- | --- | --- |
| 2000 | 90,020 | 627 | 0.70 |
| 2001 | 85,592 | 2,227 | 2.60 |
| 2002 | 88,293 | 936 | 1.06 |
| 2003 | 86,387 | 1,311 | 1.52 |
| 2004 | 51,199 | 5,870 | 11.47 |
| 2005 | 96,503 | 414 | 0.43 |
| 2006 | 96,636 | 2,892 | 2.99 |
| 2007 | 68,599 | 125 | 0.18 |
| 2008 | 103,887 | 3,544 | 3.41 |
| 2009 | 103,315 | 91 | 0.09 |
| 2010 | 81,899 | 883 | 1.08 |
| 2011 | 85,097 | 1,056 | 1.24 |
| 2012 | 87,554 | 6,017 | 6.87 |
| 2013 | 94,583 | 488 | 0.52 |
| 2014 | 86,967 | 783 | 0.90 |
| 2015 | 112,341 | 6,576 | 5.85 |
| Total | 1,418,872 | 33,840 | 2.38 |

The reference data also includes reasons for crop loss that were sporadically provided by the farmer. Fig 4 shows the numbers of barley fields affected by different drivers of crop losses. It can be seen that the main reasons for crop loss reported by farmers for barley in Finland were related to an over- or under-supply of water.

**Fig 4.**
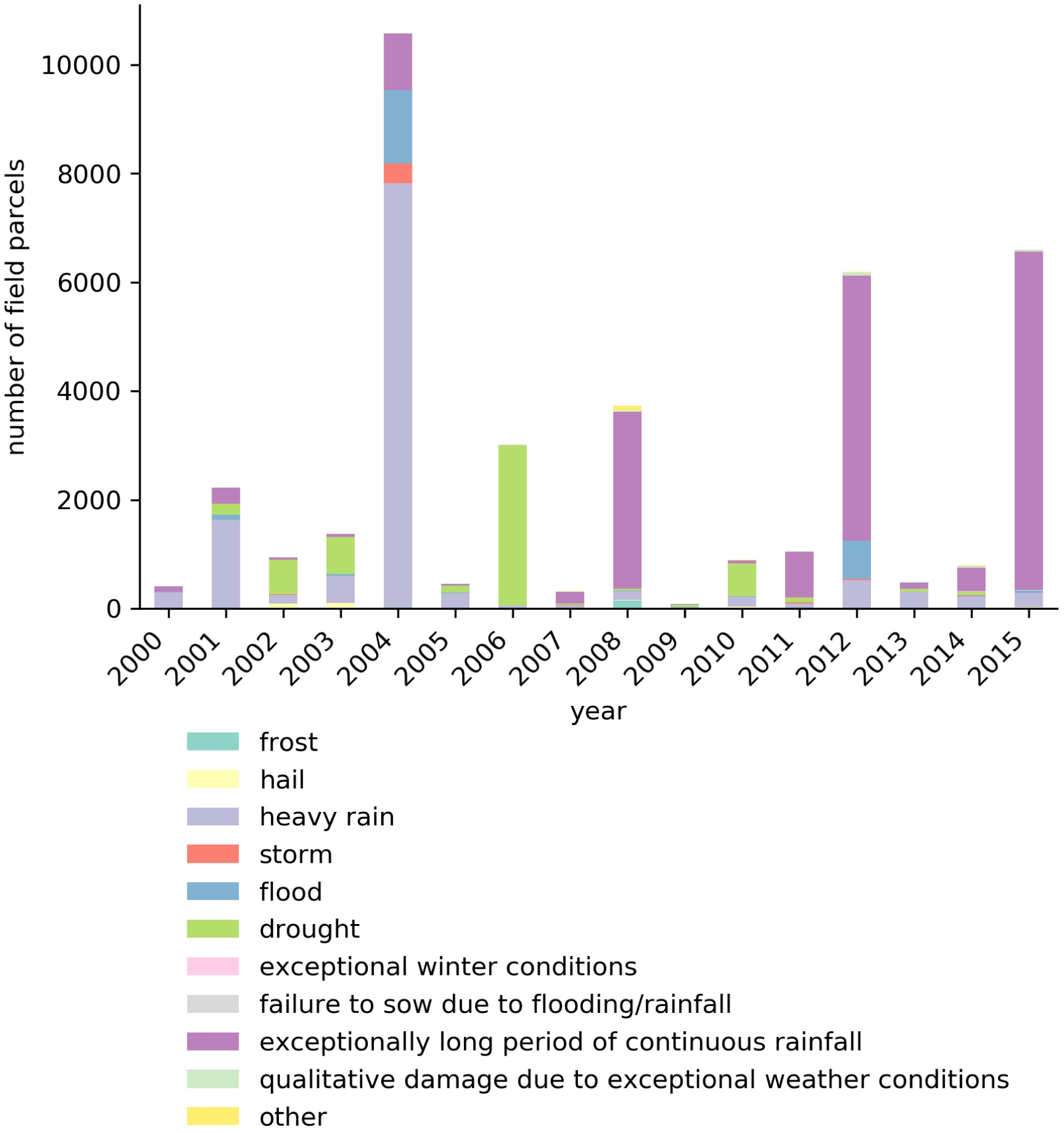
Reasons for crop loss of barley in Finland between 2000-2015. Percentage from total (33560) per type for all years: frost: 0.67%, hail: 0.79%, heavy rain: 25.70%, storm: 0.74%, flood: 5.45%, drought: 15.83%, exceptional winter conditions: 0.02%, failure to sow due to flooding or rainfall: 0.11%, exceptionally long period of continuous rainfall: 49.66%, qualitative damage caused over a large area due to exceptional weather conditions 0.53%, other: 0.50%.

### 2.2 Satellite data

Landsat 7 ETM+ (Enhanced Thematic Mapper [32]) satellite data was chosen for the study to cover the area and time frame of the reference data. Landsat 7 was launched in April 1999 and is still operating as of May 2020. It carries a multispectral sensor, which provides 8 bands covering the visible range, near-infrared and mid-infrared range as well as one thermal infrared and one panchromatic band. All bands are provided with a spatial resolution of 30 m except panchromatic and thermal infrared bands, which are provided with 15 m and 60 m resolution, respectively. The revisit time of the satellite to a specific point on earth is 16 days.

All available surface reflectance products [33] from January 2000 to December 2015 were requested from the United States Geological Survey (USGS) and downloaded using the ESPA^1^ bulk downloader. No filters were applied to the query other than the path and row indicators for the area of interest with least spatial overlap. The query resulted in 597 scenes. An overview of the number of scenes acquired per year can be found in Table 2. The surface reflectance product also includes a Quality Assessment (QA) band indicating the cloud cover based on the CFMask algorithm [34]. The QA band was used to generate a binary cloud mask. Note that, the surface reflectance product is not processed when the solar zenith angle is larger than 76 degrees. Thus data availability is limited, since the study area is above 60°North.

**Table 2.** Number of Landsat 7 ETM+ surface reflectance products acquired per year (January - December) per tile (cf. Fig 2 for location of tiles).

| year/tile | 18718 | 18918 | 19015 | 19017 | 19116 |
| --- | --- | --- | --- | --- | --- |
| 2000 | 15 | 14 | 13 | 16 | 15 |
| 2001 | 10 | 13 | 10 | 11 | 10 |
| 2002 | 11 | 13 | 10 | 11 | 12 |
| 2003 | 9 | 7 | 5 | 6 | 4 |
| 2004 | 4 | 4 | 2 | 1 | 4 |
| 2005 | 5 | 5 | 4 | 4 | 5 |
| 2006 | 5 | 4 | 4 | 5 | 2 |
| 2007 | 3 | 4 | 1 | 1 | 6 |
| 2008 | 4 | 6 | 5 | 5 | 6 |
| 2009 | 6 | 8 | 5 | 6 | 5 |
| 2010 | 4 | 5 | 4 | 3 | 4 |
| 2011 | 5 | 8 | 6 | 9 | 8 |
| 2012 | 6 | 11 | 6 | 7 | 10 |
| 2013 | 11 | 10 | 7 | 8 | 11 |
| 2014 | 7 | 8 | 11 | 11 | 9 |
| 2015 | 14 | 13 | 10 | 14 | 12 |

In 2003, Landsat 7 experienced a scan line corrector malfunction, which influenced later acquisitions by introducing gaps with missing data in the scenes. However, field parcels in Finland are much smaller than this gap, and, therefore, no correction or filling of the gaps was performed. The gaps were interpreted as missing data for each field located within the gap.

### 2.3 Data preparation

All Landsat 7 scenes were processed to create a data set in the required format to train and test the classification models, by four steps: extracting image sequences, computing NDVI time series, aggregating NDVI across time and imputing missing data. Fig 5 schematically shows these steps, which are described in detail below.

**Fig 5.**
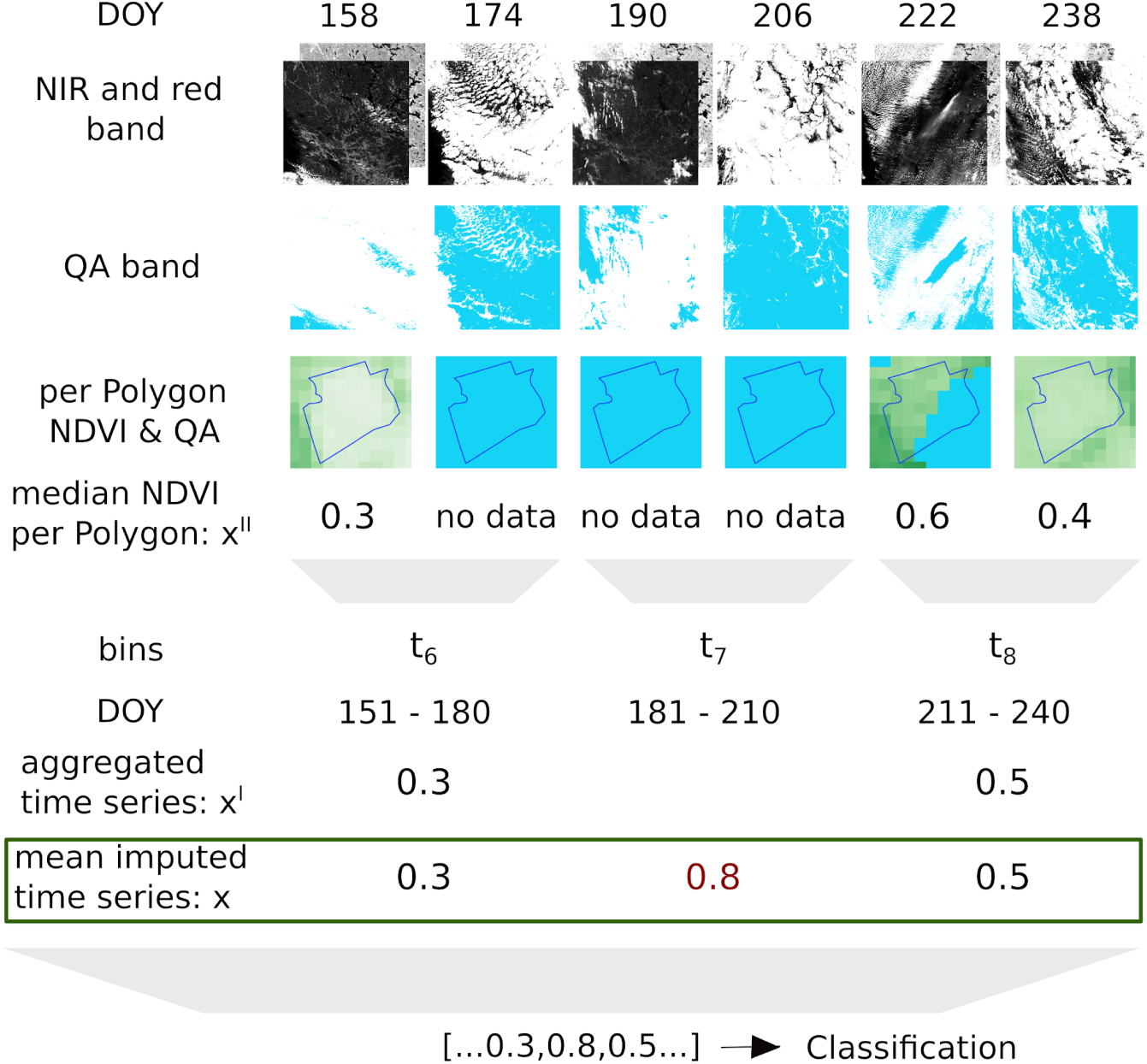
Data preparation workflow. Workflow (top to bottom) of computing time series **x** (= (0.3, 0.8, 0.5) in the green box), starting from raw Landsat 7 scenes (top) for part of a time series (DOY 158-238 2000). The 0.8 (marked in red) is the imputed mean value (of all other fields at the same time point) for t_7_. In the end, the time series is forwarded to the classification as independent features. DOY: day of year, NIR: near infrared and QA: Quality Assessment/cloud mask.

#### 2.3.1 Extracting image sequences

For each field parcel, the boundary information from the reference data was used to extract the corresponding image segments from the raster files. An image segment consists of all pixels within the field-parcel boundary. If a field parcel was in two Landsat 7 tiles, only one was kept to avoid overlap. This yielded a sequence of images (of seven bands - the six spectral bands and the pixel QA band) that were further processed to discard invalid pixels using the QA mask. These were mainly cloud pixels.

#### 2.3.2 Computing NDVI time series

The multispectral images were used to compute Normalised Difference Vegetation Index (NDVI) band according to the formula:

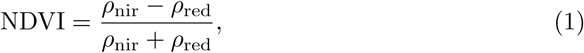

where *ρ*_nir_ and *ρ*_red_ are the pixel values of the near infrared (central wavelength 0.77 - 0.90 *µ*m) and red (central wavelength 0.63 - 0.69 *µ*m) bands, respectively. This process resulted in a sequence of NDVI images for each field parcel. From these (NDVI) image sequences, we get NDVI time series **x**″ by taking the median (NDVI) pixel.

#### 2.3.3 Aggregating NDVI across time

The temporal resolution of **x**″ refers to the frequency at which Landsat 7 scenes were captured. This is mainly a function of revisit frequency (16 days) of the satellite and cloud cover. As a result, time series **x**″ of different fields have different lengths and their time indices are not aligned. To address these two problems, we perform temporal averaging as follows: First, we form a new time scale from 1 to 365 (corresponding to each day of a year) within which each time series **x**″ is located based on the time of capture. Then, the new time scale is divided into *d* bins. The NDVI values within each bin are the mean aggregated to yield a new time series **x**′. We set *d* = 12, by which edges for 12 bins are given by *t*_1_ = [1, 30], *t*_2_ = [31, 60], *t*_3_ = [61, 90], …, *t*_12_ = [331, 360]. Some example time series **x**′ are shown in Fig 6, where red and blue lines represent fields with and without crop loss, respectively. It can be seen that, unlike a typical time series, there are many holes in **x**′. That is, even after temporal averaging the time series of each field parcel has many missing values. If we take the average of all the red and all the blue curves, then we see a general pattern of the aggregated NDVI time series for the two classes as shown in Fig 7. In each year, the top-right value shows the Pearson correlation (referred to as NDVI-corr) between the red and blue curves, indicating a high correlation between them in each year. These high correlations imply the hardness of classifying the parcels into those with loss or without loss. NDVI-corr is used later in Section 3.2 for exploring the factors to explain classification performance.

**Fig 6.**
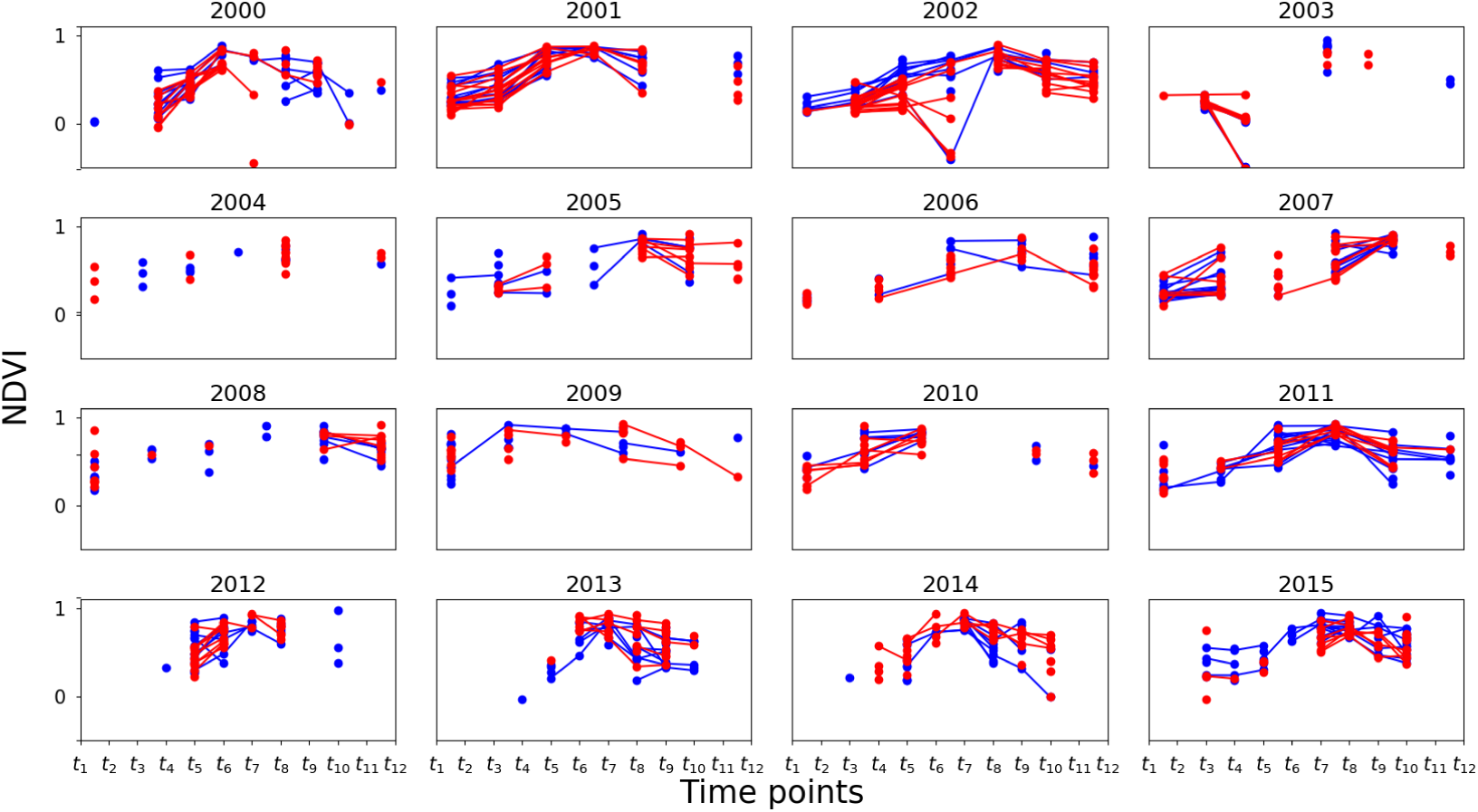
NDVI time series examples. Time series plots for some example field parcels with and without crop loss in red and blue, respectively. Note that the lack of lines between the points is due to the missing values.

**Fig 7.**
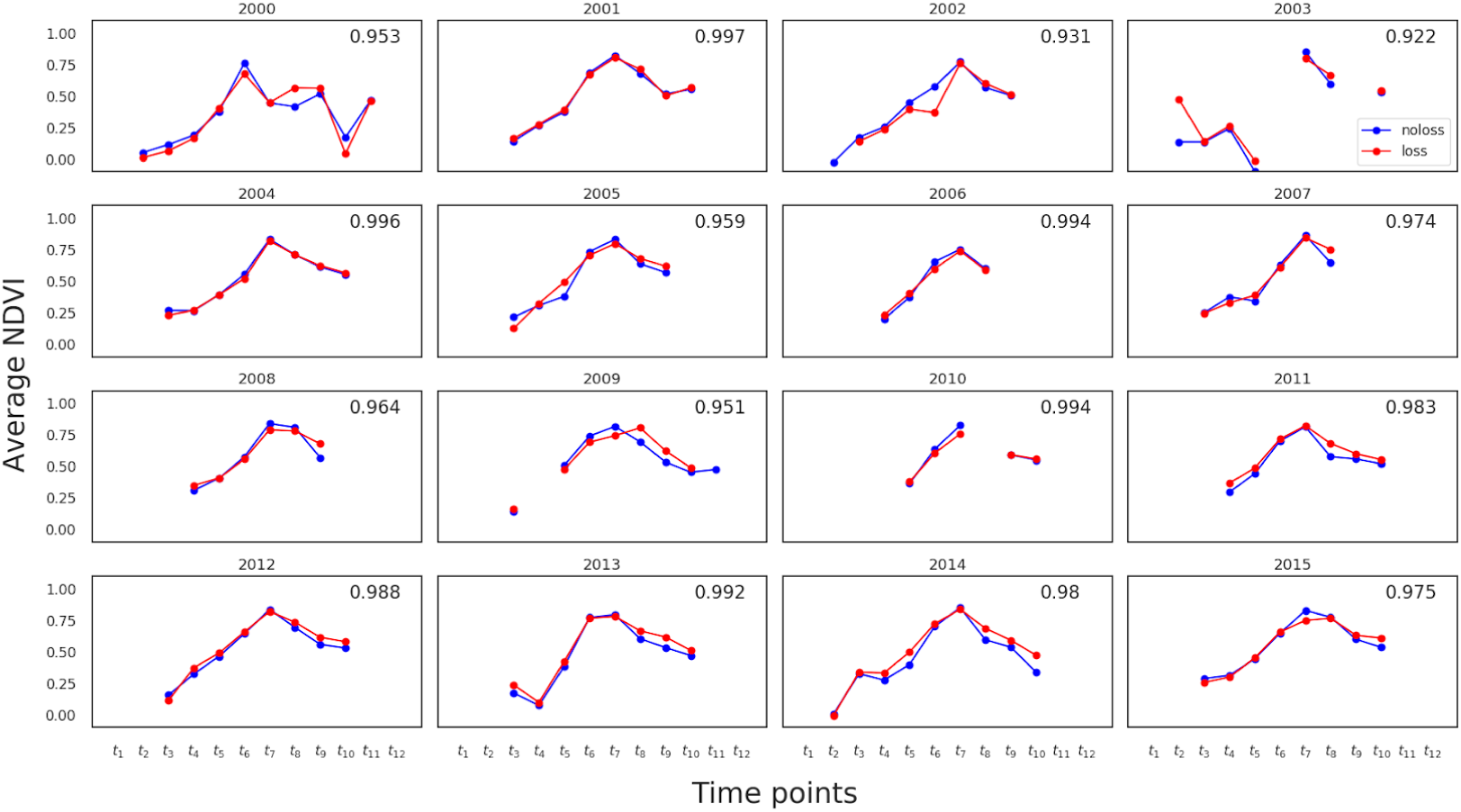
Average NDVI time series. Average NDVI time series of each year for each class (red=loss, blue=no-loss). The top-right (NDVI-corr) value in each year shows the Pearson correlation between the two curves.

#### 2.3.4 Imputing missing data

##### Missing data problem

Missing data is a common problem when dealing with satellite images due to cloud cover and other data acquisition problems. This is especially problematic for northern countries like Finland due to the low sun angle. In ideal circumstances such as no cloud cover and no acquisition problems, the time series length is approximately 22 time steps (assuming an average revisit period is 16 days for Landsat 7). However, in our case, the average length of the time series is 4 due to missing data, i.e., around 82% of the data is missing. Fig 8 shows the missing data profile for different years. In some years e.g. 2003 and 2004, the problem is more severe where more than 90% of the data are missing. We observe that for all the years, the data in the beginning and end of the year are likely to be missing. This phenomenon can be explained by the low illumination angle during winter for which no surface reflectance product is processed. For barley, these missing data points during winter should have little effect since its heading time is around beginning of July, while maturity is reached around the middle of August in this part of Finland [35]. We also see that the pattern of missing data is different for the two classes in all years, suggesting that information on the location of missing values can help improving the classification.

**Fig 8.**
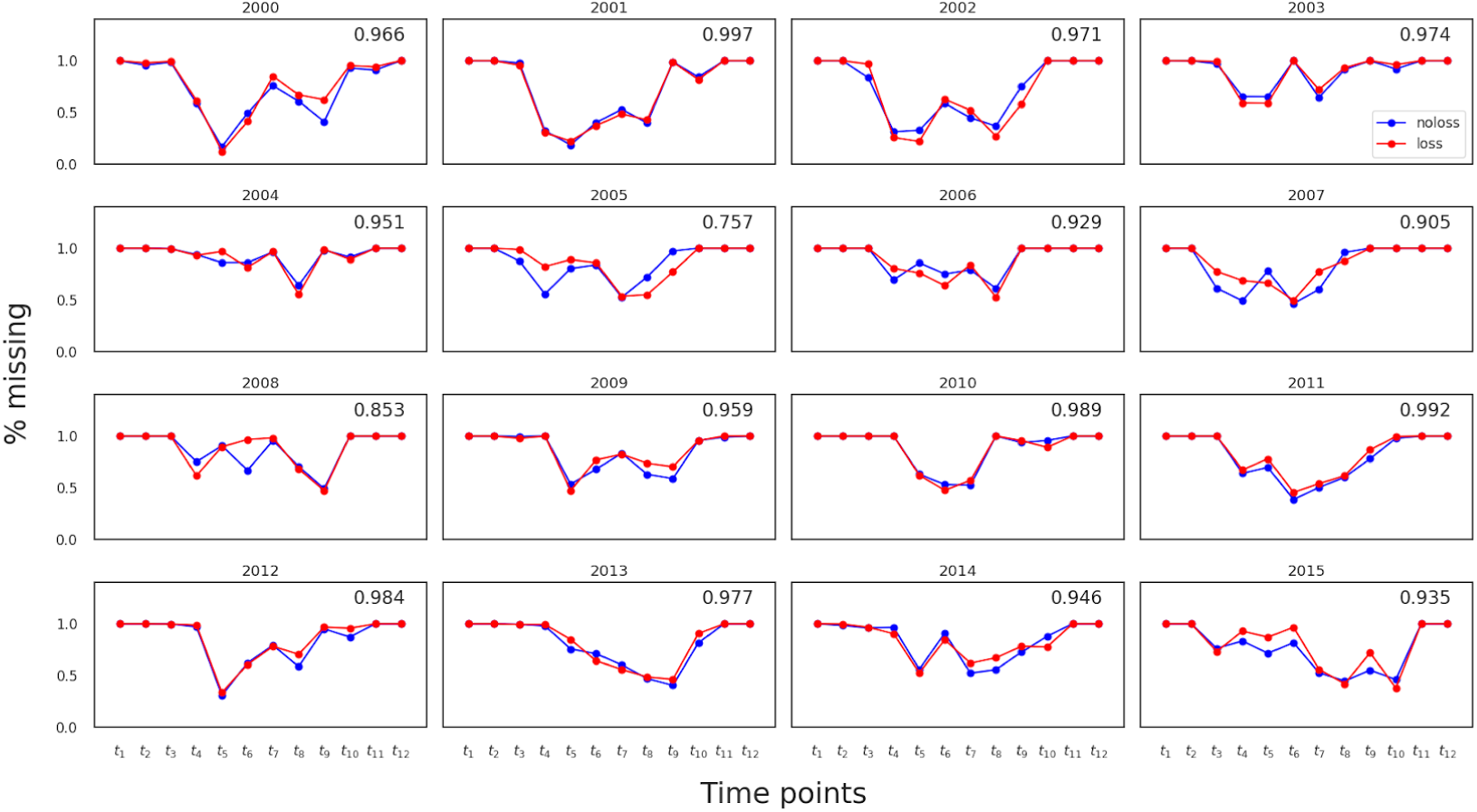
Missing data profile. Percentage of field parcels with missing values on the y-axis vs time point on the x-axis. The red and blue curves indicate crop loss and no crop loss, respectively. The top-right value in each year shows the Pearson correlation between the two curves, referred to as MD-corr (used later in Section 3.2)

##### Imputation methods for missing data

We address the missing data problem in **x**′ through mean imputation (Mean). The procedure is described using the following matrix notation for clarity. The time series **x**′ of all the fields form a matrix **X**′ where columns are the new time indices described in Section 2.3.3 and the missing entries correspond to the holes in the times series. These missing values are filled by the corresponding column mean of **X**′ yielding the matrix **X**. Each row **x**_*i*_ in the matrix **X** is the imputed time series of the field *i*.

Apart from mean imputation we also experiment with two other imputation methods: 1) missing data indicator (MI), and 2) multiple imputation by chained equation (MICE). MI generates a binary matrix **M** of the same size as the data matrix **X**, indicating the absence of a value. MICE is an iterative method which regresses each variable (column of **X**′) over the other in a round-robin fashion to compute the missing values [36]. These methods are compared in Section 3.1 to identify the best imputation strategy for classification. The imputation methods in Scikit-learn library [37] were used for the experiments.

Note that due to the severity of missing data, time series interpolation methods to fill the missing values were not considered. This is because the time series were short (average length is 4) with many instances consisting of only one or two observations. In these cases interpolation is not meaningful. Instead different imputation methods were considered to fill the missing values.

### 2.4 Classification models

Given data set 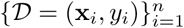 with *n* observations, the task is to learn a model *f* : **x** ↦ *y* such that *p*(*y*_*i*_) = *f* (**x**_*i*_; *θ*) where *θ* is a set of hyperparameters. We compared several classification models namely Logistic Regression (LR), Decision Trees (DT), Random Forest (RF) and Multilayer Perceptrons (MLP) [38]. Due to a large amount of missing data, which makes time series very short, we model the time points as independent features, and did not consider time series models (such as autoregressive models or recurrent neural networks) in this work. All models are implemented using Scikit-learn (version 0.22.1) Python library [37] and the details of their optimisation and model comparison is given in Section 3.1.

To compare the models, we focused on the data from year 2015 because it has the least amount of missing data (see Fig 8). Also class imbalanceness in 2015 is relatively better than other years (see Table 1). We then created a balanced data set (with 13,152 fields), with all 6,576 crop-loss fields and the equal number of no-crop-loss fields (which were sampled out of all no-crop-loss parcels by means of undersampling). This balanced data set was used for cross-validation only for the model comparison.

Before comparing the models, we first optimised the hyperparameters of each model. Table 3 shows hyperparameters that were optimised, along with their respective ranges considered and the optimal values. The results are detailed in Section 3.1

**Table 3.** Model hyperparameters, their ranges and optimal values obtained.

| Model | Parameters and Range | Optimal value |
| --- | --- | --- |
| LR | regularisation penalty = $\{1, 10\}$ | 1 |
| DT | maximum tree depth = $\{5, 10, 50\}$ | 5 |
| RF | maximum tree depth = $\{5, 10, 50\}$ | 10 |
| | #trees = $\{10, 50\}$ | 50 |
| MLP | #hidden units = $\{10, 5\}$ | 10 |

### 2.5 Performance metrics

The performance measure used to evaluate the classification models was “area under the receiver operating characteristic (ROC) curve” (AUC) [39]. AUC takes a value in the interval [0, 1], where a random classifier has a score of around 0.5 and a perfect classifier has score 1. AUC is insensitive to class imbalance which is important for this study as class imbalance is high. Further, it has a standard scale independent of the number of data points and the distribution of classes, so models trained on data from different years are directly comparable even though the number of loss and no-loss fields are different in each year. We used 10 *×* 10-fold cross-validation (CV) to compute the AUC of each model in all experiments (except in Section 3.3): In *K*-fold cross-validation first the data into *K* non-overlapping folds. Then the model is trained on *K* − 1 folds and tested on the remaining fold. This is repeated *K* times so that model is tested on each of the *K* folds. The model performance is given by average of the *K* AUC values. We used *K* = 10 and repeat 10-fold cross-validation 10 times with different random permutation of the data.

### 2.6 Within-year Classification

*Within-year classification* situation is to determine if there was a crop loss in a field (for which the crop loss information was unavailable in a year), by using fields for which crop loss data are available in the same year. We used all data of each year for cross-validation, meaning training and test parcels being from the same year. The AUC value for each year is computed using the 10×10 cross-validation procedure described in Section 2.4 to compute the AUC values for each year.

### 2.7 Between-year Classification

Collecting reference data is expensive so it would be useful to identify crop-loss fields in a year based on reference data from a different year(s). This *between-year classification* situation would be closer to future prediction of crop loss in a field, more than within-year classification. We considered two cases: single-year training and multiple-year training. For single year training case, all data from one year was used for training while all data from another year was used for testing. In the multiple year training case, test data are taken from one year and training data are taken from the remaining 15 years. For example, if test data is taken from 2015 then training data are taken from 2000 to 2014. Note that cross-validation was not used for between-year classification.

## 3 Results

### 3.1 Model comparison

We first compared machine learning models and imputation methods, to find the most appropriate model and imputation strategy to be used throughout this work. As part of identifying the optimal imputation strategy, we also decide whether or not to include indicators specifying the locations of missing values as part of the input to the model.

Table 4 shows AUCs for the different combinations of the model and imputation strategy. The method with the highest AUC was the combination of RF and Mean+MI (mean imputation and missing value indicators), followed by the pair of MLP and Mean+MI. Table 5 shows computation time of different methods, indicating the significant computational advantage of Mean and Mean+MI over MICE. Focusing on Mean+MI, we further studied model comparison over all 15 years (see Appendix 5 in detail). From this result, although RF and MLP were overall the two most accurate methods, taking computation time into account, we concluded that the combination of RF and Mean+MI is the recommended model for crop-loss classification, and used this combination for the remainder of the experiments in this study.

**Table 4.** Mean AUC of 10×10-fold CV for different models and imputation methods.

|  | Mean | Mean+MI | MICE | MICE+MI |
| --- | --- | --- | --- | --- |
| LR | 0.6903 ± 0.009 | 0.7377 ± 0.008 | 0.6473 ± 0.010 | 0.7128 ± 0.008 |
| DT | 0.7080 ± 0.008 | 0.7187 ± 0.007 | 0.6364 ± 0.010 | 0.7223 ± 0.008 |
| MLP | 0.7322 ± 0.009 | 0.7546 ± 0.008 | 0.6458 ± 0.010 | 0.7469 ± 0.008 |
| RF | 0.7514 ± 0.008 | **0.7602 ± 0.008** | 0.6515 ± 0.010 | 0.7505 ± 0.008 |

**Table 5.** Ratio of training time of four models relative to RF.

|  | Mean | Mean+MI | MICE | MICE+MI |
| --- | --- | --- | --- | --- |
| LR | 0.03 | 0.05 | 10.12 | 10.03 |
| DT | 0.04 | 0.05 | 10.36 | 10.36 |
| MLP | 12.21 | 10 | 15.91 | 20.51 |
| RF | 0.97 | **1** | 11.42 | 11.88 |

### 3.2 Within-year classification

Table 6 shows the within-year classification performance for 16 years. The average AUC across all years is 0.6884 *±* 0.027 with the best performance in 2008 with AUC = 0.795 *±* 0.008 and the worst in 2004 with AUC= 0.602 *±* 0.010. We investigated possible factors to explain the differences in those AUCs across years:

1. **NDVI correlations:** We first checked the correlation in NDVI between crop-loss and no-crop-loss fields, i.e. the correlation (NDVI-corr) between blue and red curves in Fig 7. Fig 9 shows a scatter plot of AUC against NDVI-corr for all years. We see that AUC decreases as NDVI-corr increases. For example, 2004 had the highest NDVI-corr=0.996 and the lowest AUC=0.602, whereas 2008 had the lowest NDVI-corr=0.964 and the highest AUC=0.795.
2. **Amount of missing data:** We then examined the impact of missing data on AUC. Fig 9 plots AUC against the percentage of missing data per year, where AUC decreased with increasing amount of missing data. The linear regression line, *y* = −0.3*x* + 1.0, indicates that the classification performance can improve when the amount of missing data reduces.
3. **Missing data profile correlations:** We further checked the similarity in missing data profiles of two classes by using MD-corr. Fig 9 plots AUC against MD-corr, showing that AUC decreases with MD-corr is increasing, meaning that not only the amount of missing data but also the pattern of missing data affects classification performance.

**Table 6.** Within-year classification performance (AUC) for all the years.

| Year | AUC |
| --- | --- |
| 2000 | $0.757 \pm 0.028$ |
| 2001 | $0.648 \pm 0.018$ |
| 2002 | $0.765 \pm 0.022$ |
| 2003 | $0.706 \pm 0.021$ |
| 2004 | $0.602 \pm 0.010$ |
| 2005 | $0.750 \pm 0.031$ |
| 2006 | $0.657 \pm 0.015$ |
| 2007 | $0.627 \pm 0.070$ |
| 2008 | **$0.795 \pm 0.008$** |
| 2009 | $0.647 \pm 0.070$ |
| 2010 | $0.624 \pm 0.027$ |
| 2011 | $0.673 \pm 0.023$ |
| 2012 | $0.636 \pm 0.010$ |
| 2013 | $0.679 \pm 0.033$ |
| 2014 | $0.690 \pm 0.292$ |
| 2015 | $0.755 \pm 0.008$ |
| Mean | $0.684 \pm 0.027$ |

**Fig 9.**
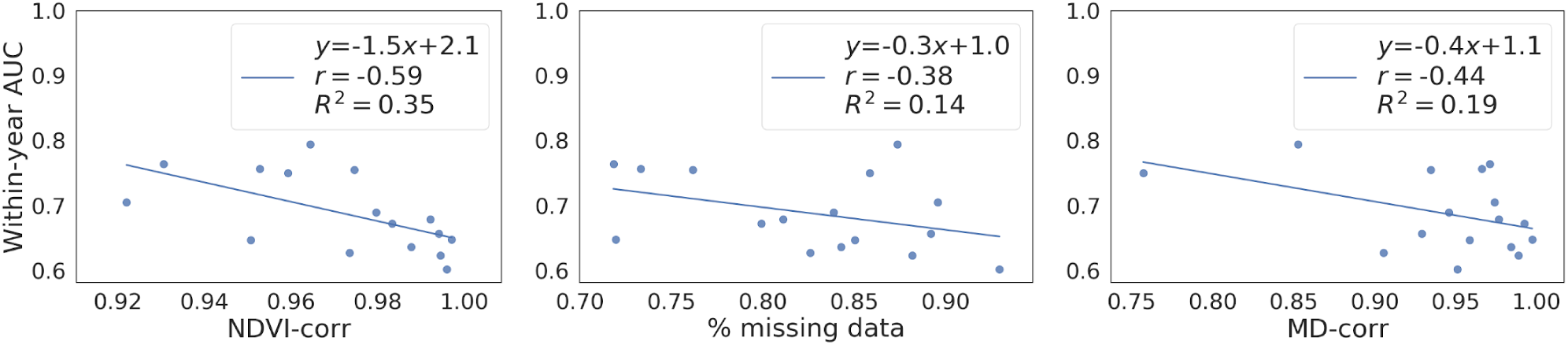
Analysis of AUC values. Effect of NDVI-corr (left), % missing data (middle) and MD-corr (right) on AUC of within-year classification. The Pearson correlation between the quantities are given by *r* in the top right corner of the each plot. The regression line and *R*^2^ are included for illustrative purpose only to highlight the inverse relationship between the quantities.

We thus can see NDVI correlation, missing data ratio and missing data profile correlation, are important factors in the data that affect classification performance. Also we hypothesise that similar missing data patterns may indirectly indicate similar weather conditions or geographical closeness of different fields, which might be useful for classification. We note that using missing data indicators as input to the classification model will be feasible in practice, as those will be available for the application at the same time as the satellite images themselves. However, missing data can be based on many reasons, such as cloud cover, data not processed to surface reflectance and scan line error so we cannot draw a causal relationship between the missingness pattern and crop-loss even when the classification accuracy is high.

#### Impact of field parcel area

We further examined the potential impact of field area on within-year classification, since larger fields are more likely to be crop-loss fields and this bias may affect classification performance. For this experiment, we focused on data from 2004, 2008, 2012 and 2015 (each had *>*3% crop-loss fields; see Table 1), to ensure that the class imbalance problem is not exacerbated when the data is divided based on the area. The field parcels are divided into three groups, depending on their area: small (*<* 1ha), medium (≥1ha and *<*3ha), and large (≥3ha). Table 7 shows the number of fields in the three groups for these four years. Table 8 shows the performance results, indicating that in each year, AUCs were approximately consistent with those obtained by using all data in Table 6 (2004: 0.602, 2008: 0.795, 2012: 0.636 and 2015: 0.755) and AUCs in different groups were close to each other. Hence, the field size would not play a significant role in the results.

**Table 7.** Ratio (%) of crop loss parcels for three groups with different areas and four years.

|  | small |  |  | medium |  |  | large |  |  |
| --- | --- | --- | --- | --- | --- | --- | --- | --- | --- |
|  | #parcels | #parcels<br>with loss | ratio<br>(%) | #parcels | #parcels<br>with loss | ratio<br>(%) | #parcels | #parcels<br>with loss | ratio<br>(%) |
| 2004 | 15,050 | 1,676 | 11.13 | 22,267 | 2,511 | 11.28 | 13,882 | 1,683 | 12.12 |
| 2008 | 30,948 | 920 | 2.97 | 43,935 | 1,444 | 3.29 | 29,004 | 1,180 | 4.07 |
| 2012 | 25,282 | 1,558 | 6.19 | 36,532 | 2,537 | 6.94 | 25,841 | 1,922 | 7.44 |
| 2015 | 31,112 | 1,548 | 4.98 | 44,998 | 2,570 | 5.71 | 36,231 | 2,458 | 6.78 |

**Table 8.** Within-year AUC of three groups with different field parcel sizes.

| Year | small | medium | large |
| --- | --- | --- | --- |
| 2004 | $0.6071 \pm 0.021$ | $0.5878 \pm 0.009$ | $0.5938 \pm 0.019$ |
| 2008 | $0.8118 \pm 0.018$ | $0.7947 \pm 0.010$ | $0.7580 \pm 0.014$ |
| 2012 | $0.6314 \pm 0.026$ | $0.6387 \pm 0.017$ | $0.6237 \pm 0.020$ |
| 2015 | $0.7605 \pm 0.023$ | $0.7600 \pm 0.010$ | $0.7386 \pm 0.015$ |

### 3.3 Between-year classification

#### Single-year training

Fig 10 visualises totally 240 AUCs of all combinations of sixteen years, by using a heat map. Many AUC values were close to 0.5, and the average AUC= 0.534 *±* 0.051. The maximum AUC was 0.665 (2003 for training and 2005 for testing) which is less than the average within-year AUC=0.688. These results indicate that training data with only one year might not be informative enough for identifying parcels with crop loss in between-year classification.

**Fig 10.**
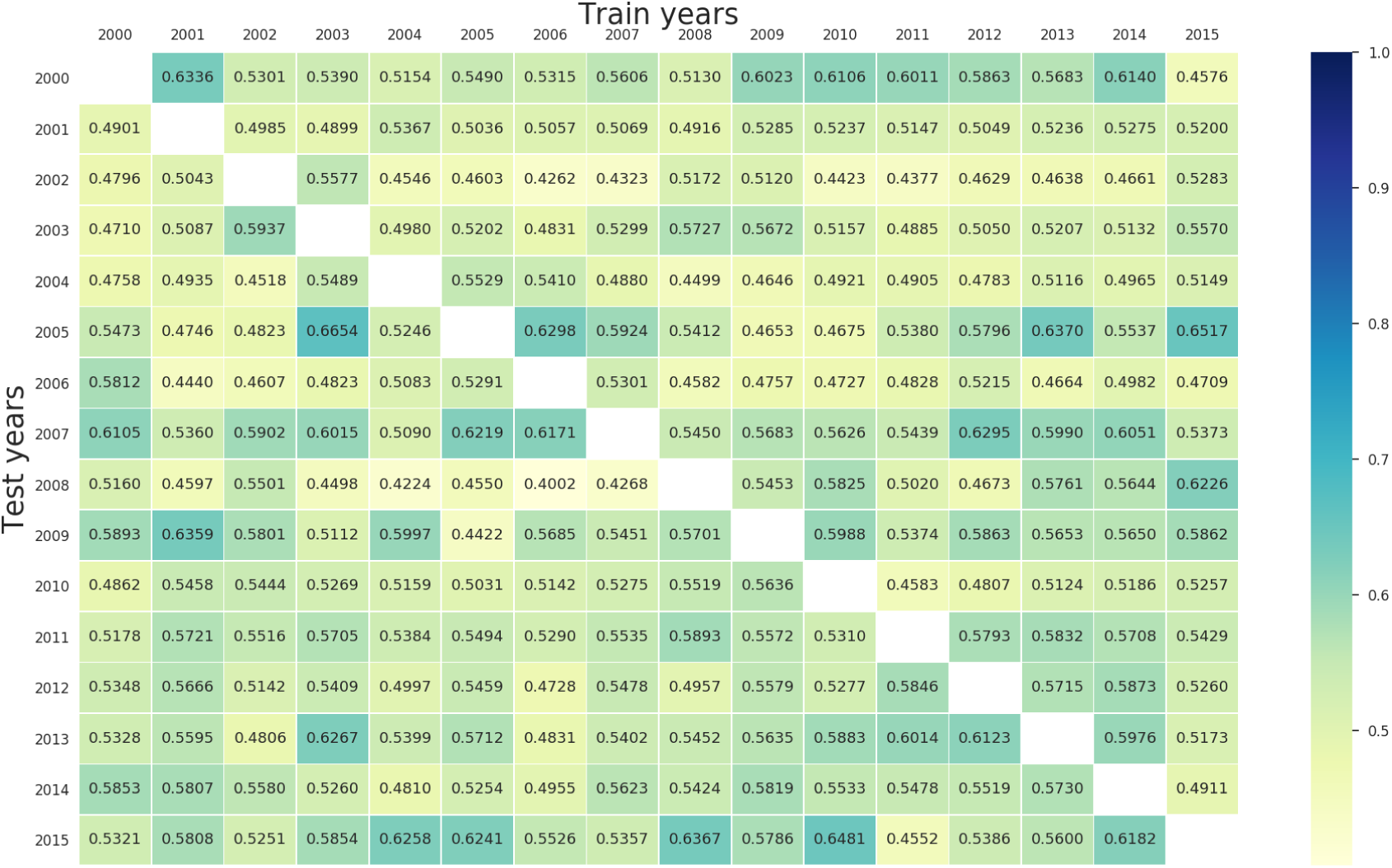
AUC values of single-year training experiment. The column and row heading indicates the year on which the model is trained and tested, respectively.

#### Multiple-year training

Table 9 shows sixteen AUC values (one for each test year) obtained by this procedure, along with the corresponding average and best AUCs of single-year training. We can see several years in which multiple-year AUC can be better than the average single-year AUC, but for all years, multiple-year AUC is always worse than the best single-year AUC. This result implies that combining data from multiple years will not improve between-year classification.

**Table 9.** Multiple year training vs Single year training (AUC).

| Year | Multiple year | Single year (average) | Single year (best) |
| --- | --- | --- | --- |
| 2000 | 0.6329 | $0.5608 \pm 0.047$ | 0.6335 |
| 2001 | 0.5226 | $0.5111 \pm 0.015$ | 0.5367 |
| 2002 | 0.4582 | $0.4764 \pm 0.039$ | 0.5576 |
| 2003 | 0.5319 | $0.5230 \pm 0.035$ | 0.5936 |
| 2004 | 0.5227 | $0.4967 \pm 0.032$ | 0.5529 |
| 2005 | 0.6287 | $0.5567 \pm 0.068$ | 0.6654 |
| 2006 | 0.4710 | $0.4921 \pm 0.036$ | 0.5811 |
| 2007 | 0.6362 | $0.5785 \pm 0.037$ | 0.6295 |
| 2008 | 0.6018 | $0.5027 \pm 0.068$ | 0.6226 |
| 2009 | 0.5703 | $0.5654 \pm 0.044$ | 0.6358 |
| 2010 | 0.5818 | $0.5184 \pm 0.028$ | 0.5636 |
| 2011 | 0.5423 | $0.5557 \pm 0.021$ | 0.5892 |
| 2012 | 0.4999 | $0.5382 \pm 0.033$ | 0.5872 |
| 2013 | 0.5548 | $0.5573 \pm 0.043$ | 0.6267 |
| 2014 | 0.5826 | $0.5437 \pm 0.033$ | 0.5853 |
| 2015 | 0.6093 | $0.5731 \pm 0.052$ | 0.6480 |

## 4 Discussion

We have trained machine learning models to classify field parcels with and without crop loss, using NDVI values derived from Landsat 7 data. Several models were compared and tested in two different scenarios, namely within-year and between-year classification. The results showed that within-year classification is highly possible, while between-year classification is still hard. The resulting models have many applications, for example, they can be used by insurance companies or government agencies for verifying crop-loss claims. However, their performance will be contingent upon the availability of good reference data that captures all possible variations in environmental and biophysical conditions.

Our study showed that a major challenge to improve the classification performance is the amount of missing data. Figure 9 shows the possibility of achieving high AUC if the missing data ratio was low. In our case more than 80% of data are missing and despite that the average within-year AUC=0.688 with the possibility to increasing up to 90% when the problem is not severe. The issue might be improved by using newer spaceborne multispectral data, such as those from Sentinel 2 which has higher temporal and spatial resolution that could mitigate the effect of missing data. Furthermore, Sentinel 2 data can be combined with RADAR data from Sentinel 1 to get a denser time series and avoid occlusion from clouds to provide more detailed information about the growing pattern of agricultural fields. Obviously, such data are available only from the most recent years, and therefore cannot be combined with our crop loss data, which had covered many years but until 2015.

Between-year classification allows identification of crop-loss fields without any reference data in the same year so improving its performance would be a good future target. Our result in Section 3.3 implies that NDVI data alone might not be sufficient for improving the performance of between-year classification. Incorporating temperature and precipitation data along with the crop-loss reasons can give a more complete picture of the factors affecting the crop loss. Based on the promising results of within-year classification, augmenting high resolution satellite data with weather variables would be useful in achieving high performance for between-year classification.

Another direction in which our work could be extended is the use of more flexible machine learning models. For example, convolutional neural networks could use the full image as input and recurrent neural networks could explicitly model the time dependency. These models have the potential to increase the classification performance. However, these models require a large amount of data to train properly, and cannot be directly applied to our data, which has a huge amount of missing data and consists of relatively short time series. The machine learning models in our study are rather simpler as they are more robust against the limitations in our data.

## 5 Conclusion

We have proved the feasibility of training a machine learning model to identify crop loss at field parcel scale using NDVI data derived from Landsat 7 satellite images. Experiments across sixteen years showed that field parcels from a given year can be classified into those with and without crop loss when the model can be trained on data from the same year. However, the ability to classify parcels from other years is limited. Missing data, which occupied more than 80% of our satellite images, deteriorated the classification performance. Preliminary analysis indicated that within-year classification performance can be improved if the missing data ratio was reduced. As long as we can assume that the NDVI times series of barley crop is similar with minor variation due to environmental and biophysical conditions then we believe that findings of this study can be generalised to barley fields in other countries.

## Acknowledgments

This joint research between Aalto University and Natural Resources Institute Finland (Luke) is funded by the AIPSE programme of Academy of Finland through the AI-CropPro project; decision number 315896 (Aalto) and 316172 (Luke).

## Appendix

### Model comparison for all years

In Section 3.1, we performed model comparison on 2015 data to determine the best model. Table 10 extends these results to the other 15 years. Here all models were implemented with Mean+MI imputation strategy. We see that for all years, RF and MLP have have similar AUC values and outperform DT and LR. However, MLP was less robust that RF, i.e., it did not converge even after 500 iterations for several years (2005, 2007, 2009, 2011, 2013, 2014) and took longer to train (Table 5). Thus, considering performance, stability and training time, we selected RF for further experiments.

**Table 10.** AUC of different models with Mean+MI imputation strategy.

| Year | RF | MLP | DT | LR |
| --- | --- | --- | --- | --- |
| 2000 | **0.7643 ± 0.029** | 0.7471 ± 0.037 | 0.6662 ± 0.044 | 0.7139 ± 0.046 |
| 2001 | **0.6509 ± 0.021** | 0.6409 ± 0.018 | 0.5735 ± 0.018 | 0.6223 ± 0.023 |
| 2002 | 0.7555 ± 0.036 | **0.7582 ± 0.025** | 0.6771 ± 0.053 | 0.7051 ± 0.031 |
| 2003 | **0.6954 ± 0.032** | 0.6779 ± 0.029 | 0.6602 ± 0.028 | 0.6625 ± 0.028 |
| 2004 | 0.5920 ± 0.015 | **0.5943 ± 0.015** | 0.5721 ± 0.012 | 0.5849 ± 0.007 |
| 2005 | **0.7699 ± 0.034** | 0.7552 ± 0.030 | 0.6935 ± 0.041 | 0.7452 ± 0.038 |
| 2006 | 0.6624 ± 0.021 | **0.6632 ± 0.024** | 0.6176 ± 0.026 | 0.6392 ± 0.025 |
| 2007 | 0.6844 ± 0.097 | **0.7048 ± 0.101** | 0.6029 ± 0.105 | 0.6824 ± 0.088 |
| 2008 | 0.7840 ± 0.012 | **0.7871 ± 0.012** | 0.7474 ± 0.012 | 0.7667 ± 0.016 |
| 2009 | 0.7146 ± 0.133 | **0.7548 ± 0.087** | 0.6242 ± 0.124 | 0.7046 ± 0.084 |
| 2010 | 0.6418 ± 0.031 | **0.6549 ± 0.036** | 0.6085 ± 0.024 | 0.6478 ± 0.041 |
| 2011 | **0.6839 ± 0.054** | 0.6708 ± 0.058 | 0.6333 ± 0.049 | 0.6624 ± 0.050 |
| 2012 | **0.6360 ± 0.016** | 0.6344 ± 0.014 | 0.5985 ± 0.015 | 0.6169 ± 0.020 |
| 2013 | **0.6771 ± 0.050** | 0.6729 ± 0.049 | 0.6354 ± 0.052 | 0.6888 ± 0.060 |
| 2014 | **0.7213 ± 0.030** | 0.7103 ± 0.035 | 0.6315 ± 0.046 | 0.6959 ± 0.032 |

### Visualising within-year classification results

Figure 11 shows the observed and predicted crop loss of 1000 field parcels for 2008. We choose 2008 because it has 1) ‘sufficient’ number of crop-loss parcels to divide into train and test sets, 2) relatively low missing data and 3) highest AUC. The following procedure was used to generate the plot. First data was randomly split into two sets: train and test. The test set consisted of 500 random crop-loss and 500 random no-crop-loss field parcels. All the remaining data formed the train set that was used to train the model (combination of RF and Mean+MI). Then the ROC curve of the trained model was analysed and the ‘optimal’ classification threshold determined to be the one that maximises the sensitivity and specificity of the model (also called Youden’s J-index [40]). Finally, the trained model and the classification threshold was used to classify the field parcels in the test set. It should be noted that Youden’s J-index is just one way to determine the optimal operating point on the ROC curve. An optimal point of a classifier is a trade-off between the cost of detecting false positives against the cost of failing to detect positive which is always application dependent.

**Fig 11.**
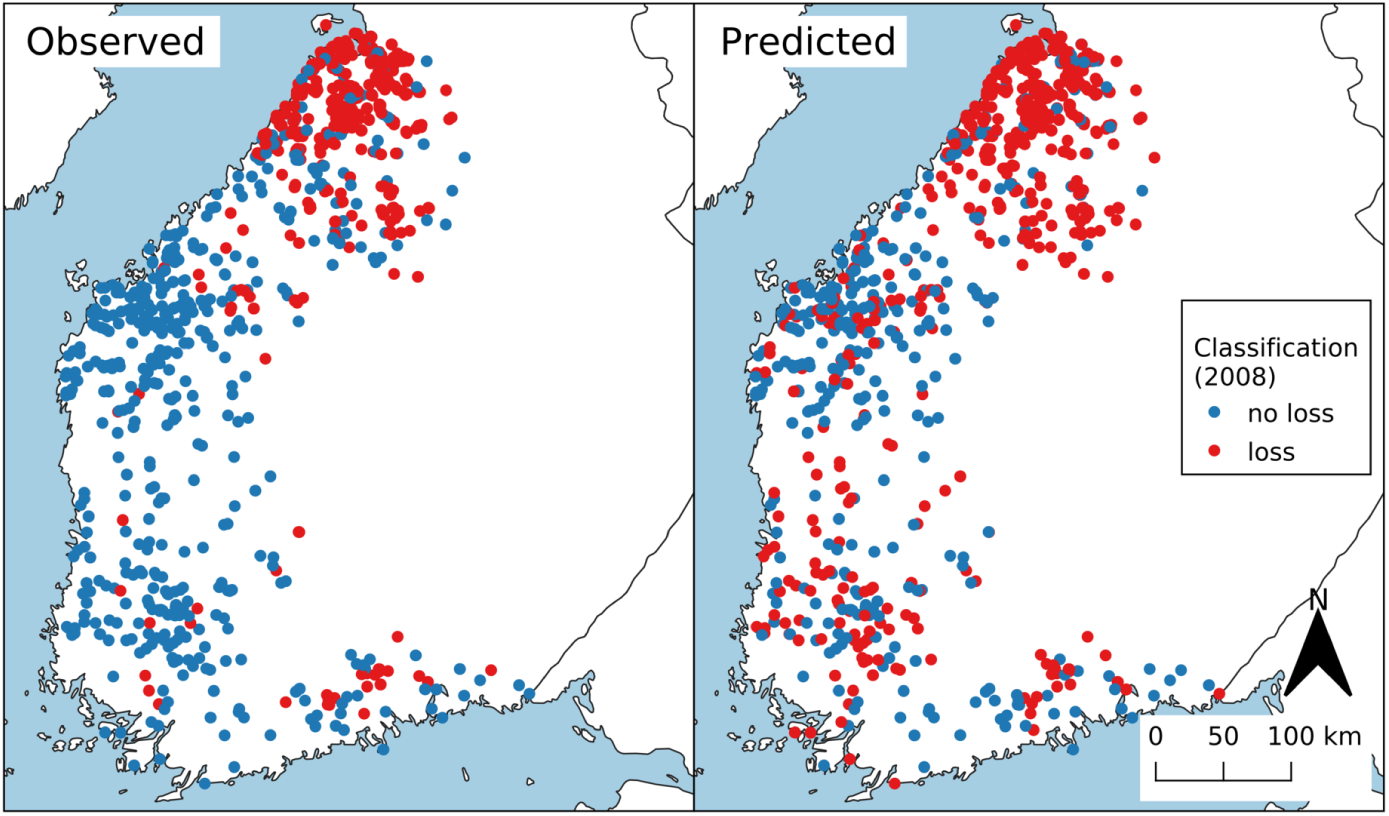
Classification result. Observed (left) vs predicted (right) crop loss of 1000 randomly chosen field parcels in 2008 simplified as points. Red point indicates croploss of the fieldparcel, blue point indicates no loss in 2008.

## Footnotes

1 ESPA stands for EROS Science Processing Architecture, and EROS stands for Earth Resources Observation and Science.

